# Analysis of Microsatellite Instability Intensity in Single-Cell Resolution with scMnT reveals Tumor Heterogeneity in Colorectal Cancer

**DOI:** 10.1101/2025.05.09.653227

**Authors:** Gyumin Park, Jihwan Park

## Abstract

Although microsatellite instability (MSI) is traditionally regarded as a binary trait, growing evidence indicates that it exists on a continuum, with higher intensity of MSI correlating with better prognosis. However, accurate quantification of MSI intensity is challenging due to the heterogeneous cellular composition of the tumor microenvironment. To this end, we introduce scMnT, a bioinformatic method for identifying MSI cells at single-cell resolution and quantifying MSI intensity based on microsatellite allele lengths. Applying scMnT to scRNA-seq datasets produced robust results and revealed substantial variation in MSI intensity among colorectal cancer patients, with higher MSI intensity correlating with improved responses to immunotherapy. Transcriptomic analyses revealed that MSI intensity-low cancer cells were enriched in goblet signatures, whereas MSI intensity-high cancer cells exhibited strong stem cell characteristics, suggesting a potential link between MSI intensity and cancer cell origin. Our approach provides a novel framework for analyzing MSI at the single-cell level, allowing new opportunities for MSI research.

## Main

MSI is a genetic hypermutation phenotype characterized by an increased mutation rate in the microsatellite regions due to DNA mismatch repair (MMR) deficiency and is relatively frequent in gastric, breast, endometrial, cervical, and especially colorectal cancers (CRC) [1, 2]. Notably, MSI tumors are known to respond well to immune checkpoint blockade (ICB) therapy, underscoring MSI as a critical biomarker for therapeutic screening [3–5]. While MSI is often regarded as a binary phenotype, studies have shown that MSI can be measured as a quantitative phenotype, where higher MSI intensity has been associated with increased tumor-infiltrating T lymphocytes and better response to PD-1 blockade [6, 7]. These previous studies highlight the importance of quantifying MSI intensity, rather than merely detecting MSI, to better stratify patients eligible for ICB therapy.

Despite growing recognition of MSI as a quantitative phenotype, current detection methods fall short in this regard. PCR-based MSI detection is limited by the small number of loci assessed, while immunohistochemistry (IHC) only detects the presence or absence of MMR proteins. Although next-generation sequencing (NGS)-based tools such as MSIsensor-pro provide an MSI score (calculated as the percentage of unstable loci) they do not account for the presence of non-malignant cells (e.g., normal cells) nor do they distinguish the varying MSI intensity between cancer cells [8–11]. Among several possible improvements, selectively analyzing tumor cells using single-cell resolution techniques could enhance the quantification of MSI intensity. Indeed, single-cell RNA sequencing (scRNA-seq) has become indispensable for cancer research, enabling the study of tumor heterogeneity at unprecedented resolution [12, 13]. A recently introduced tool, MSIsensor-RNA, enables detection of MSI from various NGS data including scRNA-seq data [14]. While it achieves excellent accuracy, it infers MSI status indirectly through the gene expression pattern of MSI CRC signature genes and does not assess MSI status or intensity at single-cell resolution. This results in immune cells and stromal cells of MSI tumors exhibiting higher MSI scores compared to such cells of MSS tumors. Thus, the lack of tools for single-cell resolution MSI analysis highlights the need for new approaches [15].

Here, we introduce scMnT, a bioinformatic method for analyzing tumor MSI status from scRNA-seq data. scMnT builds on our previously published microsatellite analysis software, NanoMnT, which accounts for the error profile of Oxford Nanopore sequencing platforms during microsatellite allele calling [16]. scMnT leverages microsatellite length information from reads mapped to microsatellite regions and groups them by their cell barcodes, enabling the identification of MSI cells at single-cell resolution and precise quantification of MSI intensity. Analysis of publicly available CRC datasets [17, 18] revealed both inter- and intra-tumoral variation in MSI intensity as well as transcriptomic properties associated with MSI intensity. Our work offers a novel method for quantifying MSI intensity and new opportunities analyzing MSI cancer cells at the single-cell level. scMnT is publicly available at GitHub: https://github.com/18parkky/NanoMnT/tree/scMnT.

## Results

### Inference of MSI status in single-cell resolution

An overview of scMnT is provided in Figure 1a (see Methods). First, sequencing reads mapped to mononucleotide-repeat microsatellite loci within the genome are collected and assigned to individual cells based on their cell barcodes, enabling the reconstruction of microsatellite profiles at single-cell resolution. Next, the relative microsatellite allele size is measured using the GRCh38 reference allele as a surrogate for the host genotype because genotype information is typically unavailable from scRNA-seq data alone. While this approach does not provide locus-specific genotyping, we have previously observed that it is sufficient for inferring MSI status from sequencing data [16]. Subsequently, the average and standard deviation of relative allele sizes for each cell are calculated. MSI cells typically exhibit decreased average allele sizes and increased variability (higher standard deviation), reflecting the accumulation of deletion mutations in mononucleotide repeats [19]. In contrast, microsatellite stable (MSS) cells maintain allele size distributions that are similar to the reference genome, with low average and low standard deviation values. Finally, the MSI score is calculated as the product of the average and standard deviation, multiplied by –1, integrating the two metrics into a single composite score. These scores are then used to infer the MSI status of individual cells or user-defined cell clusters. For the latter, scMnT performs this inference by statistically comparing the MSI score distribution of a target cluster (e.g., cancer cells) to that of normal, MSS cells (e.g., immune cells). The comparison is conducted using the Kolmogorov-Smirnov test, and the effect size is reported using Cliff’s delta. Clusters with a statistically significant p-value (e.g., <0.01) and effect size (e.g., >0.4) are considered MSI.

**Figure 1.**
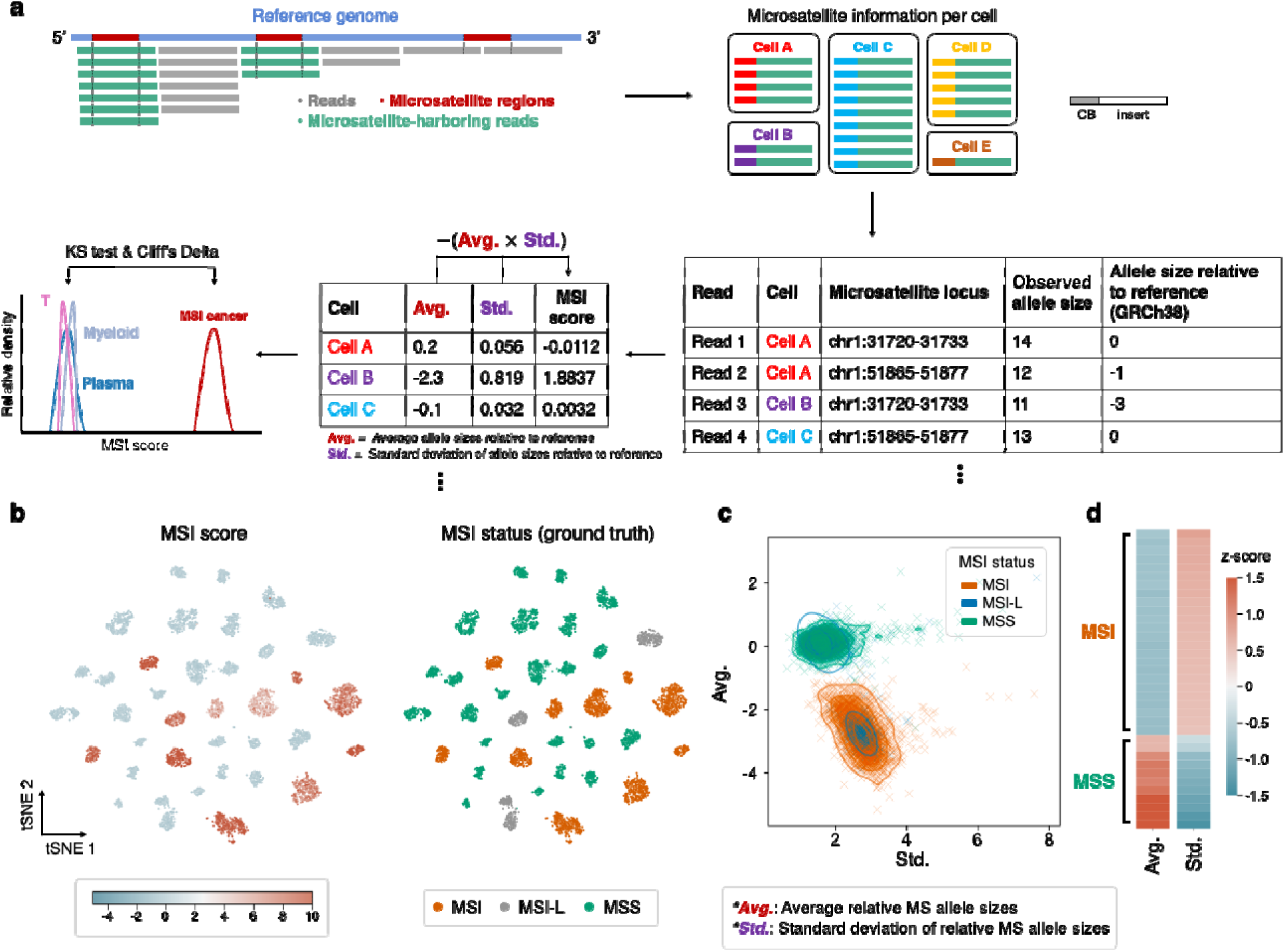
Overview of scMnT and its application to cancer cell line data. **a**, Schematic diagram depicting the workflow of scMnT. From the BAM file generated by Cell Ranger, sequencing reads that map to mononucleotide-repeat microsatellite regions are grouped by cell barcode (CB). The allele size (i.e., length of the microsatellite) relative to the reference genome (GRCh38) i determined for each read, which is used to calculate the MSI score for each cell. Using user-provided cell type annotations, the MSI scores can be compared among cell types using the Kolmogorov-Smirnov (KS) test and Cliff’s Delta values. **b**, t-SNE plot of cancer cells from a scRNA-seq dataset from Kinker et al. (2020) showing accurate MSI identification results from scMnT. Ground-truth MSI status for each cell line was obtained from Cellosaurus and literatures. **c**, Scatter plot of cancer cells based on the standard deviation of their relative microsatellite allele sizes (x-axis) and their average relative microsatellite allele sizes (y-axis), demonstrating that both metrics contribute to describing the MSI status of cells. Each ‘x’ represents a cell. **d**, Heatmap of each cancer cell line colored by their average-(left) and their standard deviation-(right) of the relative microsatellite allele sizes. Each bar represents a cancer cell line.

Applying scMnT to a publicly available cancer cell line scRNA-seq dataset [20] produced robust results, with MSI scores for each cell line aligning well with their known MSI status (Fig. 1b, Extended Data Fig. 1a, Supplementary Table 1). As expected, we found that both the average and standard deviation values provide valuable information about the MSI status of single cells (Fig. 1c). MSI cell lines exhibited varying degrees of MSI intensity, whereas MSS cell lines maintained relatively stable MSI intensity near zero, indicating that the observed variability of MSI scores among MSI cell lines likely represent true biological differences (Extended Data Fig. 1b). Notably, cancer cell lines known to be MSI-L, which is clinically considered to be the same as MSS [1], exhibited varying MSI scores that evidently resembled MSI scores of either MSI or MSS.

While users can test MSI in user-defined clusters of interest, the MSI status of individual cells can also be inferred directly from their MSI scores. In such cases, using a threshold of 1.5 to distinguish MSI from MSS cells yielded the best performance in terms of sensitivity and specificity (Extended Data Fig. 1c). However, it is important to note that the optimal threshold may vary depending on the single-cell chemistry and sequencing instrument, as these factors determine the number of PCR cycles applied to the cDNA molecules, which are known to introduce significant amount of noise known as PCR stutter [21, 22]. To ensure that the number of detected microsatellite loci per cell does not lead to bias in MSI score, we inspected the relationship between the two metrics (Extended Data Fig. 1d). Regardless of the number of loci detected, MSS cells consistently exhibited low MSI scores. In contrast, MSI cells showed a weak but significant correlation between MSI score and the number of detected loci. Nonetheless, this effect did not impact the overall MSI intensity of cell lines, indicating that MSI classification remains robust (Extended Data Fig. 1e).

### Variations in MSI intensity among CRC patients

We applied our approach to a scRNA-seq dataset of 10 CRC patients from the CRC-SG1 cohort and KUL cohort [18]. UMAP visualization of cells after preprocessing with Scanpy [23] (Methods) revealed that all cells except for subsets of epithelial cells displayed low MSI scores (Fig. 2a, Extended Data Fig. 2a). Sub-clustering of epithelial cells revealed multiple patient-specific clusters—representing the malignant cells—as well as a single cluster containing cells from multiple patients, which represent the normal intestinal epithelial cells. Again, substantial variation in MSI intensity was observed among the MSI patients, but not in MSS patients, further supporting the quantitative properties of MSI phenotype (Fig. 2b). Furthermore, MSI scores were markedly higher in longer microsatellites, as expected given that increased length is known to promote strand slippage. Notably, normal cells and MSS cancer cells also exhibited elevated MSI scores in microsatellites longer than 20 bp, likely due to PCR stutter artifacts, which are similarly correlated with microsatellite length.

**Figure 2.**
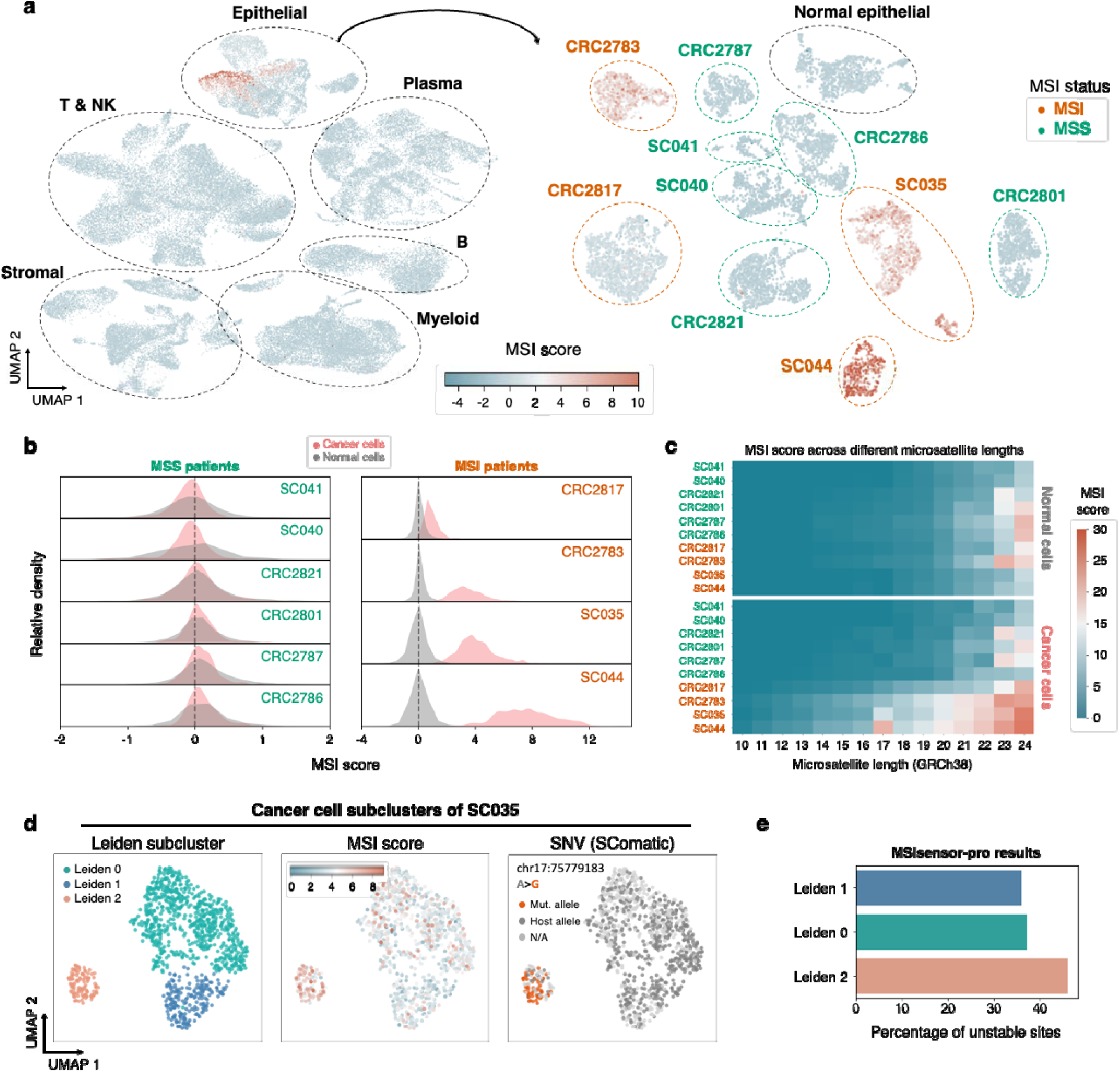
Inter- and intra-tumoral heterogeneity in MSI intensity across colorectal cancer patients. **a**, UMAP visualization of all cells (left) and epithelial cells (right) from a colorectal cancer (CRC) patient dataset from Joanito et al. (2022). Only cells with at least 10 reads mapped to microsatellite regions are shown, and cells are colored by their MSI scores. **b**, Kernel density estimate (KDE) plot of MSI scores of normal cells (gray) and cancer cells (red). **c**, Heatmap showing the average MSI scores across different microsatellite lengths in normal cells (top) and cancer cells (bottom). **d**, UMAP visualization of cancer cells from patient SC035, colored by their Leiden cluster (left), MSI scores (middle), and SNV (right). **e**, MSIsensor-pro results of each SC035 cancer cell Leiden cluster.

In patient SC035, cancer cells initially formed two distinct clusters in the UMAP embedding (Fig. 2a), prompting a more detailed analysis by sub-clustering the cancer cells separately. A refined UMAP embedding revealed three distinct cancer cell clusters with varying MSI intensities (Fig. 2d, Extended Data Fig. 2b). Suspecting that these clusters may reflect polyclonal origins of CRC, we analyzed their single-nucleotide variants (SNVs) using SComatic [24]. Hierarchical clustering based on SNV features further supported the hypothesis that these clusters arose from distinct clones (Methods, Extended Data Fig. 2c, 2d). Moreover, we verified that this observation was not an artifact by applying MSIsensor-pro (using pseudo-bulks from each SC035 cancer subcluster as inputs), which produced results (Fig. 2e) concordant with those from scMnT (Extended Data Fig. 2e). These results suggest that intratumoral heterogeneity may extend to variations in MSI intensity.

### Transcriptomic signatures correlated with MSI intensity

While high MSI intensity is associated with favorable prognosis and improved response to ICB treatments [6, 7], whether MSI intensity is driven by biological factors or determined randomly remains unclear. To investigate this, we sought transcriptomic signatures correlated with MSI intensity. We expanded the number of MSI patients for analysis by incorporating an additional comprehensive CRC scRNA-seq dataset from a recent study by Chen et al. [17], which consists of 10 MSI patients and includes patient metadata on ICB response and tumor regression ratios (Extended Data Fig. 3a, 3b).

Once more, we observed substantial variability in MSI intensity among CRC patients (Extended Data Fig. 3c). Patients were classified into MSI-LI, MSI-MI, and MSI-HI (not to be confused with traditional definition of MSI-L/MSI-H) based on their MSI intensity (Methods, Extended Data Fig. 3d, Supplementary Table 2), and differential gene expression analysis was performed between MSI-LI and MSI-HI cancer cells (Fig. 3a). Genes enriched for MSI-LI cancer cells included well known markers of goblet cells (e.g., *MUC2*, *ATOH1*) and enterocytes (e.g., *TFF1*, *PIGR*), whereas genes enriched for MSI-HI cancer cells were associated with intestinal stem cell signature (e.g., *ASCL2*, *RGMB*, *SOX4*, *LEF1*) (Extended Data Fig. 3e).

**Figure 3.**
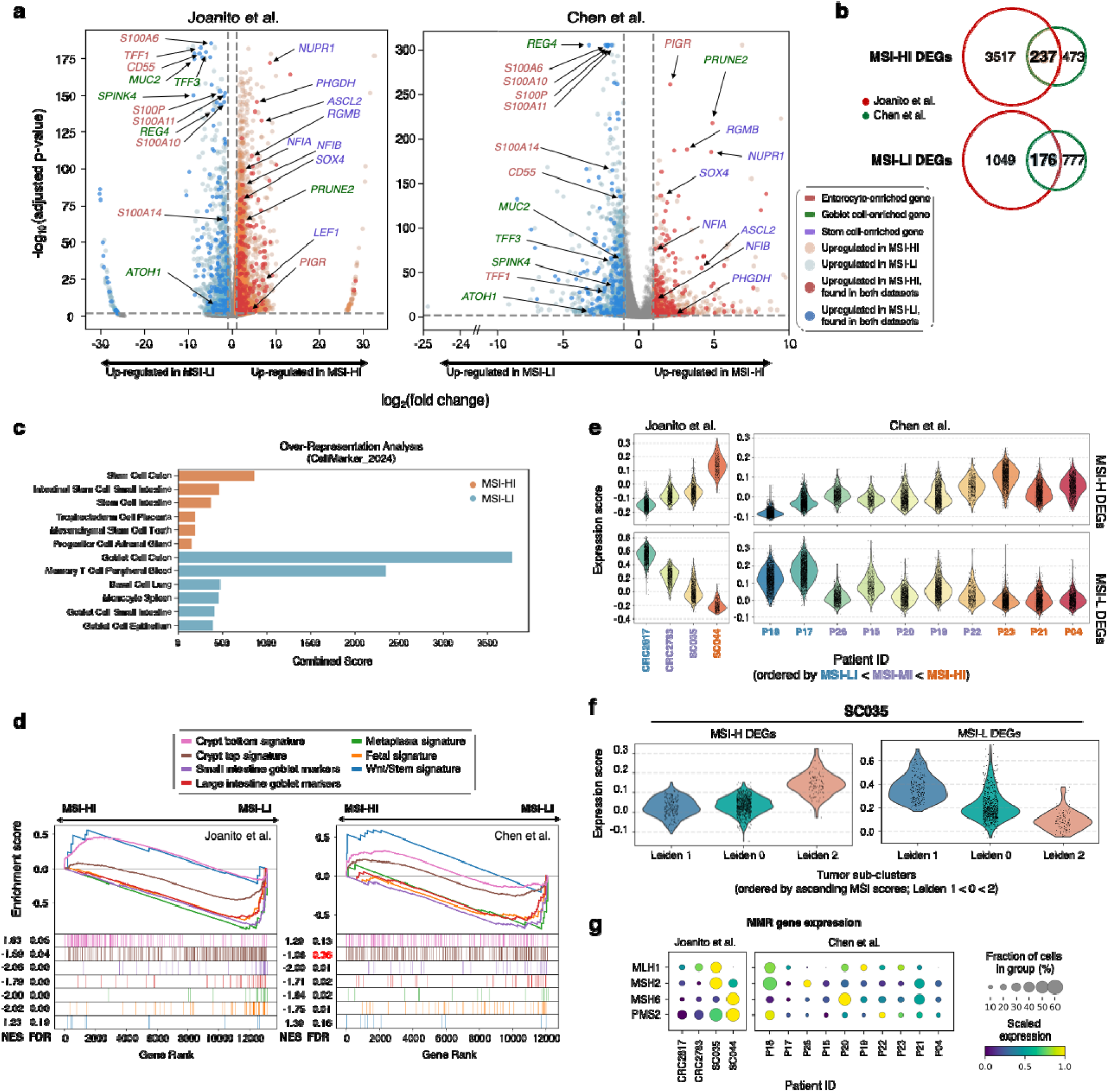
Tumor-intrinsic transcriptomic signatures associated with MSI intensity. **a**, Volcano plot displaying DEGs in MSI-LI and MSI-HI cancer cells from the Joanito et al. (2022) dataset (left) and Chen et al. (2024) dataset (right). **b**, Venn diagram showing the number of significant DEGs identified in cancer cells from the two datasets. **c**, Bar plot visualizing the overrepresentation analysis (ORA) results for DEGs identified in cancer cells from both datasets. **d**, Gene set enrichment analysis (GSEA) visualization of DEGs identified in cancer cells from both datasets, using gene sets from the literature. **e**, Violin plot showing the average expression of MSI-HI and MSI-LI-associated DEGs in cancer cells across all patients. **f**, Violin plot showing the average expression of MSI-HI and MSI-LI-associated DEGs in cancer cells of patient SC035. **g**, Dot plot visualizing the expression of MMR genes in cancer cells across all patients.

Using genes that were identified as differentially expressed genes (DEGs) in both datasets (Fig. 3b, Supplementary Table 3) as inputs, over-representation analysis (ORA) produced similar results, suggesting that cancer origin may be a possible factor that determines MSI intensity (Fig. 3c, Methods). We also noticed that MSI-LI cancer cells were characterized by high expression of S100 family genes, which are associated with fetal signatures [25, 26]. Given that CRC arising from sessile serrated lesions were found to be strongly associated with gastric metaplasia and fetal signatures [27], we performed gene set enrichment analysis (GSEA) to test for relevant pathways (Fig. 3d, Methods). Interestingly, both metaplasia, fetal signatures and crypt top signatures were enriched in MSI-LI cancer cells, while Wnt/stem and crypt bottom signatures were enriched in MSI-HI cancer cells.

Next, we calculated module scores for MSI-HI DEGs and MSI-LI DEGs across cancer cells from individual patients and found a positive relationship between MSI-HI DEG module scores and MSI intensity, and an inverse relationship for MSI-LI DEG module scores (Fig. 3e). Notably, a similar pattern was observed within the cancer subclusters of patient SC035 (Fig. 3f). Additionally, we examined the expression of MLH1 and other MMR genes in cancer cells, as MLH1 loss—often due to promoter hypermethylation—is a hallmark of sporadic MSI CRC [28–30]. Strikingly, cancer cells of several patients displayed mRNA expression of MLH1, whereas patients with the highest MSI intensity (SC044 and P04) did not (Fig. 3g). Other important MMR genes were detectable across all patients.

### High MSI intensity is associated with better ICB response and higher baseline T cell abundance

In agreement with previous reports [6, 7], we also observed that patients who achieved complete response (CR) showed higher MSI scores than patients who had partial response (PR) (Fig. 4a). Furthermore, tumor regression ratios were also correlated with the patient MSI scores, though the robustness of correlation analysis was limited by the number of patients available.

**Figure 4.**
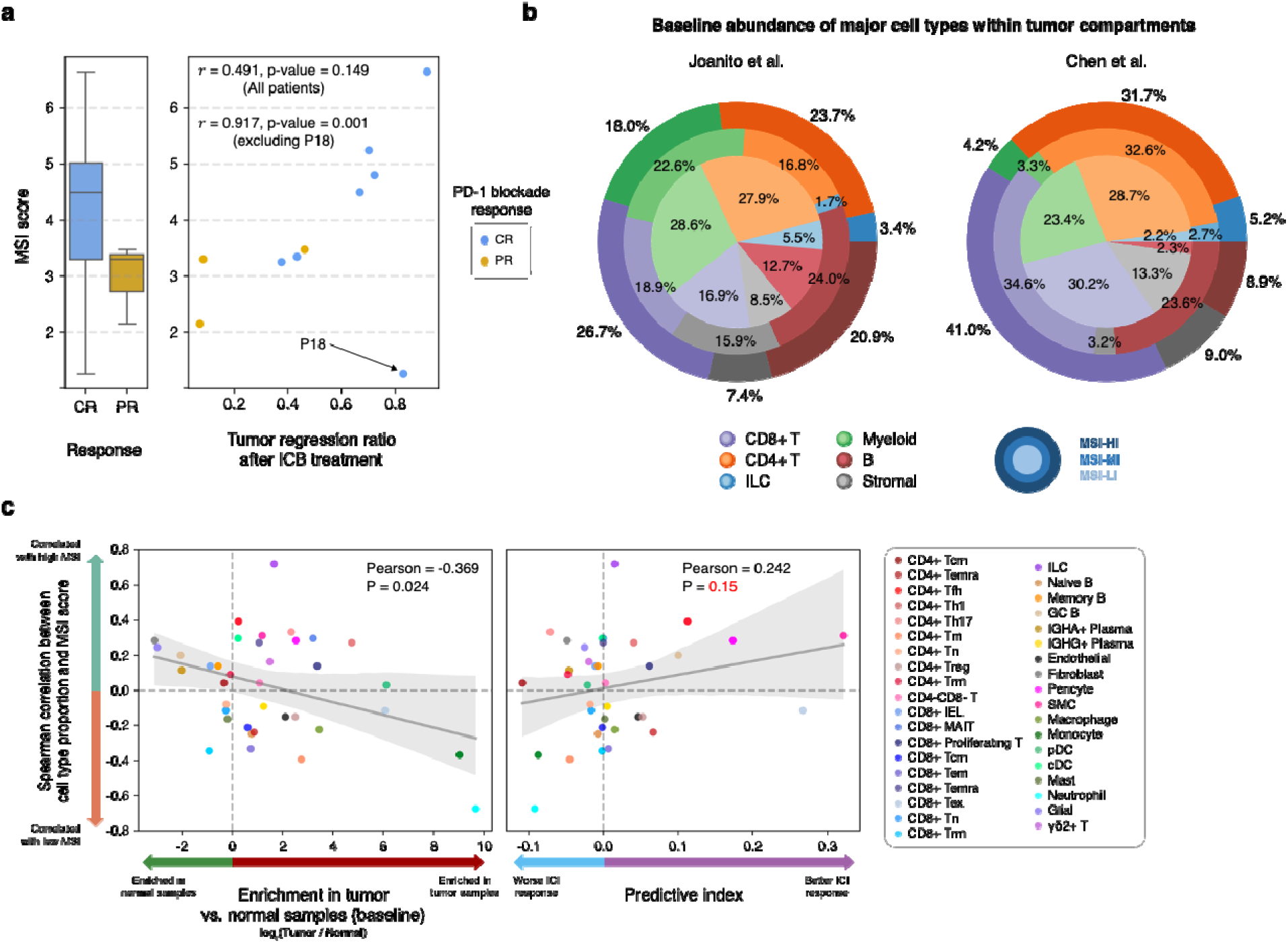
Association of MSI score with response to immune checkpoint blockade therapy and baseline abundance of various cell types. **a**, Association of MSI score and response to ICB therapy. Box plot comparing the distribution of MSI scores (of cancer cells) between patients who displayed complete response versus patients who displayed partial response (left) and scatter plot displaying the relationship between the tumor regression ratio following ICB therapy and MSI score of each patient (right). **b**, Comparison of baseline abundance of major cell types within tumor environments between MSI-LI (inner), MSI-MI (middle), and MSI-HI (outer) patients. **c**, Scatter plots showing, for each cell type, the Spearman correlation between its proportion and the MSI score (y-axis) against two variables: sample enrichment (left, x-axis) and predictive index (right, x-axis).

Examination of baseline cellular abundance of various cell types within the tumor microenvironment (TME) across the three MSI intensity classes revealed cell types whose abundance correlated with MSI intensity (Fig. 4b). In both datasets, CD8+ T cells were more abundant in tumors with higher MSI intensity, following the order of MSI-HI > MSI-MI > MSI-LI, suggesting that elevated immune infiltration may underlie the improved ICB response seen in MSI-HI CRC. Additionally, MSI-HI and MSI-MI tumors had higher baseline proportions of B cells compared to MSI-LI tumors, consistent with previous findings linking B cell presence to enhanced immunotherapy responses [31–34]. Conversely, MSI-LI tumors displayed a greater abundance of myeloid cells relative to MSI-HI and MSI-MI tumors. No other cell types showed obvious associations with MSI intensity.

To further investigate these relationships, we leveraged the high-resolution cell-type annotations from Chen et al. [17] to explore associations between baseline TME composition and MSI intensity at a finer granularity (Fig. 4c). Tumor-enriched cell types (e.g., CD8+ Tex, CD4+ Treg, endothelial cells) were modestly enriched in tumors with lower MSI intensities, whereas normal tissue-enriched cell types (e.g., GC B, Memory B) were enriched in tumors with higher MSI intensities. Additionally, cell types enriched in tumors with higher MSI intensities exhibited a higher predictive index, which reflects the correlation between baseline cellular proportions and tumor size changes following ICB treatment [34]. This suggests that higher MSI intensities are associated with a TME favorable to ICB response, although the association did not reach statistical significance (p=0.15).

## Discussion

In this study, we introduced scMnT, which is designed to (1) identify MSI cells at single-cell resolution and (2) quantify MSI intensity by analyzing the mean and standard deviation of microsatellite allele lengths. Using scRNA-seq datasets of CRC cohorts, we validated the robustness of our method by accurately distinguishing MSI cancer cells from MSS cells and quantitatively measuring the MSI intensity of individual cells. These results demonstrate that scMnT reliably detects MSI cells at single-cell resolution, enabling analysis of heterogeneous samples containing both MSI and MSS cells. scMnT also clarifies the MSI status of ‘MSI-L’ cancer cells, which are clinically classified as MSS, enabling definitive assessment of their microsatellite instability at the single-cell level. Our method also addresses a key challenge in scRNA-seq studies: annotating cancer cells. Unlike normal cell types, cancer cells often lack well-defined marker genes [35], complicating their identification—especially for MSI cancers, which typically do not exhibit common genomic features such as copy number variations [36]. scMnT thus provides a complementary strategy to improve cancer cell annotation in single-cell datasets. In summary, scMnT offers a robust and scalable framework for MSI detection in single-cell transcriptomic data, enhancing our ability to study intratumoral heterogeneity and cancer cell identity in MSI-driven tumors.

We also performed a series of analyses to elucidate biological factors associated with MSI intensity. Through comparative analyses of MSI-LI and MSI-HI cancer cells, we found that MSI-LI tumors exhibited strong fetal, metaplastic, crypt-bottom, and goblet cell signatures, whereas MSI-HI tumors were enriched for stem, Wnt, and crypt top signatures. These results suggest that MSI intensity may be shaped by biological factors rather than arising purely from stochastic processes. A previous comprehensive study reported that CRCs originating from differentiated intestinal epithelial cells (enterocytes and goblet cells) display strong gastric metaplasia signatures, whereas those arising from intestinal stem cells are characterized by prominent Wnt signaling [27]. In light of this context, our results may indicate a potential link between the cell of origin and MSI intensity.

As others have reported, we corroborate the existence of intertumoral heterogeneity in MSI intensity among CRC patients and its correlation with ICB treatment response. However, notable variations persist. For instance, patient 18 from the Chen et al. dataset exhibited a high tumor regression ratio despite showing the lowest MSI intensity (Fig. 4a). It is important to recognize that therapeutic outcomes are influenced by multiple factors beyond MSI intensity, including HLA genotype [37, 38], baseline immune infiltration [17, 39], defects in antigen presentation machinery [40, 41], Wnt pathway activation [42], presence of immunosuppressive cells [43, 44], and the degree of intratumoral heterogeneity [45, 46]. Nonetheless, patients with high MSI intensity generally exhibit better responses to ICB treatment and therefore should be prioritized for ICB treatment. This is of particular clinical relevance since around half of MSI patients fail to achieve a durable response to ICB treatment [45]. A more nuanced, quantitative understanding of MSI intensity — rather than relying on a binary classification — may help improve patient selection for immunotherapy. Furthermore, we emphasize that MSI intensity is not synonymous with tumor mutational burden (TMB). Although both metrics reflect the somatic mutational load, MSI intensity specifically captures indel-driven frameshift mutations, which are more likely to produce novel immunogenic neoantigens. In contrast, TMB accounts for both single nucleotide variants (SNVs) and indels, with SNVs often generating mutations less likely to elicit strong immune responses.

A particularly intriguing observation from our analysis was the detectable mRNA expression of MLH1 in MSI cancer cells of several MSI patients (Fig. 3g). While loss-of-function or hypomorphic mutations in mismatch repair (MMR) genes may explain this observation, such mutations are relatively uncommon in sporadic MSI colorectal cancer (CRC). Instead, most cases arise from epigenetic silencing of the MLH1 promoter via hypermethylation [47–49]. Thus, the presence of MLH1 transcripts suggests that MMR deficiency may not be a fixed state, but rather a reversible or heterogeneous phenotype. This raises the possibility that the duration or timing of MMR deficiency could influence accumulation of MSI events and therefore drive the intertumoral variation of MSI intensity of MSI tumors.

We acknowledge several limitations of this study. scMnT requires the raw files (e.g., FASTQ or BAM) and relies on a small subset of reads that align to microsatellite regions within intragenic areas, which may not be consistently captured across different cancer types. Additionally, although we have proposed several biological hypotheses — including a potential link between MSI intensity and cell of origin, as well as the possibility of MMR deficiency reversion — these hypotheses require experimental validation and further analysis in a larger cohort. Importantly, we were unable to define an MSI score threshold for selecting patients eligible for ICB therapy — a critical and long-standing clinical need. As mentioned above, this threshold is likely to vary depending on sequencing protocols (e.g., library preparation chemistry and sequencing platform) and cancer type. Future studies should therefore aim to standardize sequencing parameters within each cancer type to establish robust MSI score thresholds. We anticipate that scMnT will serve as a foundation for future studies investigating the biological relevance of MSI heterogeneity and its impact on cancer evolution and immunotherapy response.

## Methods

### scRNA-seq data preprocessing

For the SG1 and KUL cohort of CRC patients from Joanito et al. [18], raw data were downloaded from the European Genome-phenome Archive (EGA) database under accession numbers <u>EGAD00001008555</u>, <u>EGAD00001008584</u>, and <u>EGAD00001008585</u>.

For the CRC patient cohort from Chen et al. [17], raw data were downloaded from the Genome Sequence Archive under accession number <u>PRJCA018173</u>. Gene expression matrices were generated from the raw data using 10x Genomics Cell Ranger v8.0.1 [50], and author-provided cell type annotations were overlaid onto the expression matrices. Droplets with doublet score > μ + σ, computed using Scrublet [51], were considered doublets and discarded. Furthermore, droplets with cell barcodes not present in the authors’ annotations were excluded, as they had not passed quality control in the original studies.

The remaining droplets were processed using the Scanpy package [23] as follows: (1) droplets with fewer than 300 detected genes and genes expressed in fewer than 10 cells were removed; (2) UMI counts were normalized using `scanpy.pp.normalize_total` and log-transformed with `scanpy.pp.log1p`; (3) 2,000 highly variable genes, selected using `scanpy.pp.highly_variable_function`, were used for principal component analysis (PCA); and (4) 40 PCA components were used for UMAP with `scanpy.pp.neighbors` and `scanpy.pp.umap` [52, 53]. For the cancer cell line dataset from Kinker et al. [20], which employed a custom multiplexing strategy, the gene expression matrix was downloaded from the Gene Expression Omnibus (GEO) (accession number <u>GSE157220</u>) and was used for subsequent analyses.

### Reconstruction of single-cell microsatellite allele profiles

Microsatellite allele information was summarized at the read level using the `getAlleleTable` command from NanoMnT v1.0.0, which requires a list of target microsatellite regions and a BAM file as input. A- and T-repeat mononucleotide microsatellite regions ranging from >9 to <24 nucleotides in length were identified in the GRCh38 reference genome using Krait v1.3.3 [54]. BAM files tagged with cell barcodes and unique molecular identifiers generated by Cell Ranger (`possorted_genome_bam.bam`) were used as input for NanoMnT. Reads containing indel errors within the flanking regions of microsatellites were discarded. The remaining reads were grouped by cell barcodes, and for each cell, the average and standard deviation of the relative microsatellite allele sizes were calculated. These cell-level values were then overlaid onto the preprocessed Scanpy objects for downstream analyses.

### SNV analysis using SComatic

Precomputed cell type annotations and Cell Ranger output BAM files from tumor samples of patient SC035 were prepared as inputs for SComatic. The following SComatic scripts were executed in order: `SplitBamCellTypes.py`, `BaseCellCounter.py`, `MergeBaseCellCounts.py`, `BaseCellCalling.step1.py`, `BaseCellCalling.step2.py`. SNVs that passed all filters (‘Multiple_cell_types’, ‘LC_Upstream’, ‘Clustered’, ‘LC_Downstream’, ‘Min_cell_types’, ‘Cell_type_noise’, ‘Noisy_site’, ‘Multi-allelic’) were then processed using SComatic script `SingleCellGenotype.py`. SNVs supported by fewer than three reads or represented in fewer than 100 cells were discarded. The zygosity of each SNV locus was assessed by examining alleles reported by normal cells. Monoallelic loci (i.e., all normal cells reporting a single allele) were selected for downstream analysis. To compare the allele frequencies across the four cell groups in patient SC035 – normal cells and tumor cells from Leiden clusters 0-2 (as shown in Fig. 2d-e)– only SNVs represented in at least five cells per group were retained, resulting in 134 SNV loci. Hierarchical clustering was performed using the `clustermap` function from the Python package Seaborn [55], which internally uses Scipy’s `cluster.hierarchy.linkage` function [56]. Finally, SNVs whose alternate allele frequency exceeded 0.1 in any of the tumor Leiden cluster were visualized (Extended Data Fig. 2d). Allele frequencies for all four groups across each SNV locus are provided in Supplementary Table 4.

### Classification of patients based on their MSI intensity

MSI CRC patients were classified into three groups—MSI-LI, MSI-MI, or MSI-HI—based on the z-score standardized average MSI scores of their cancer cells. Patients with average scores below −0.5 were classified as MSI-LI, those above 0.5 as MSI-HI, and those with scores between −0.5 and 0.5 as MSI-MI. The average score of each patient is available in Supplementary Table 2.

### Identification of DEGs

Differentially expressed genes (DEGs) between MSI-LI and MSI-HI groups were identified using Scanpy’s `rank_genes_groups` function with the Wilcoxon method. Genes with a log fold change > 1 or < −1 and an adjusted p-value ≤ 0.01 were considered enriched in MSI-HI or MSI-LI, respectively. Genes meeting these criteria in both the Joanito et al. and Chen et al. datasets (Fig. 3b) were selected for further analysis.

### ORA and GSEA analysis

Both over-representation analysis (ORA) and gene set enrichment analysis (GSEA) were performed using the GSEApy package [57]. For ORA, we used EnrichR [58] with CellMarker_2024 [59] as the reference database to identify enriched cell-type markers. GSEA was conducted using curated gene sets derived from two studies [18, 60] and CellMarker 2024 (Supplementary Table 5).

### Characterization of DEGs using Human Protein Atlas data

The expression of selected DEGs identified in both datasets (Fig. 3a) was visualized across enterocytes, goblet cells, and intestinal stem cells using the Human Protein Atlas ‘Single Cell Type’ dataset [61]. For each gene, normalized transcript per million (nTPM) values were aggregated by cell type and converted into relative proportions to assess its expression distribution across the three cell types.

### Regression and statistical analysis

Linear regression analysis and statistical testing (Spearman correlation, Pearson correlation, and Kolmogorov-Smirnov test) has been performed using the Python package Seaborn (v0.13.2).

### Data visualization

All data visualizations have been performed using the Python packages Seaborn and Matplotlib (v3.9.2) and were adjusted using Microsoft PowerPoints.

## Data availability

All data presented in this study are publicly available in the European Genome-phenome Archive (EGA) database (accession number EGAD00001008555 and EGAD00001008585), Genome Sequence Archive (accession number PRJCA08173), and Gene Expression Omnibus (GEO) (accession number GSE157220).

## Code availability

Codes for scMnT and tutorials for running scMnT were deposited at GitHub and are publicly available (https://github.com/18parkky/NanoMnT/tree/scMnT). Codes used for the analyses in this study were also deposited at GitHub and are available (https://github.com/18parkky/CRC_MSI_intensity_analysis).

## Acknowledgements

We thank the authors of the two studies for granting access to the raw sequencing data from their cohorts. This study was supported by the National Research Foundation of Korea (NRF) (RS-2024-00335026, RS-2024-00403622), Korea Technology and Information Promotion Agency for SMEs (RS-2024-00506966), and a grant of the Korea-US Collaborative Research Fund (KUCRF), funded by the Ministry of Science and ICT and Ministry of Health & Welfare, Republic of Korea (RS-2024-00466906)

## Contributions

Both G.P. and J.P. conceptualized the study and prepared the original manuscript. G.P. designed and implemented scMnT, led the formal analysis and visualization. J.P. supervised the study and reviewed the manuscript.

## Ethics declarations

The authors declare no competing interests.

---

**Extended Data Figure 1.**
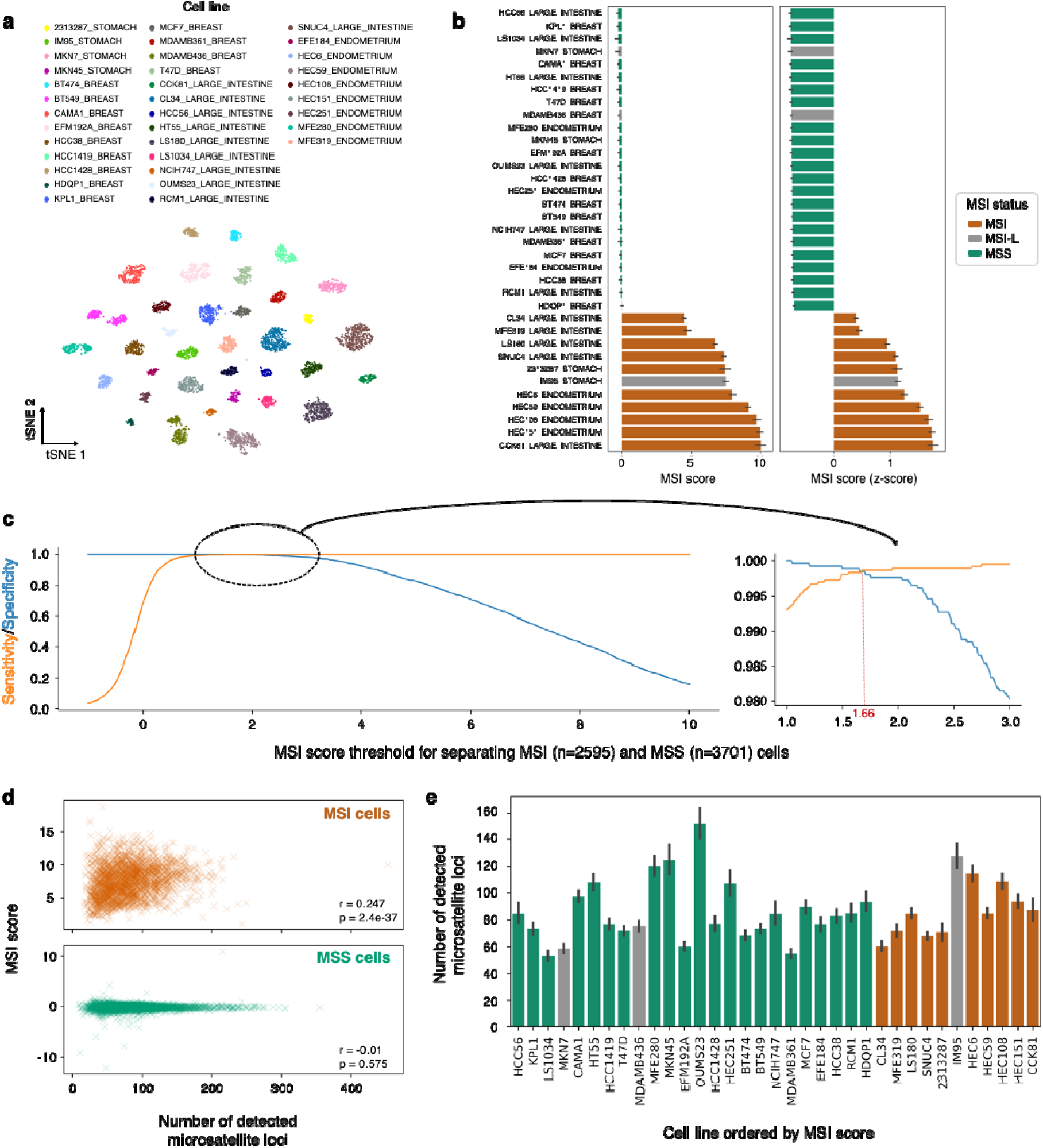
scMnT provides robust identification of MSI cancer cells from scRNA-seq data and demonstrates high specificity and sensitivity. **a**, UMAP visualization of cancer cells, colored by cell line name. **b**, Bar plot visualizing the MSI score (left) and standardized MSI scores (right) of each cell line, colored by their known MSI status. **c**, Line plot showing the specificity (blue) and sensitivity (yellow) of scMnT in identifying MSI cells across different MSI score thresholds. Specificity and sensitivity are equal at a threshold of 1.66. **d**. Scatter plots showing the relationship between the number of detected microsatellite loci and the MSI score of each cell. Each ‘x’ represents a single cell. Pearson correlation coefficient (r) and p-value (p) are shown in each plot. **e**, Bar plot showing the average number of detected microsatellite loci across different cell lines. Cell lines are ordered in ascending order based on their MSI scores.

**Extended Data Figure 2.**
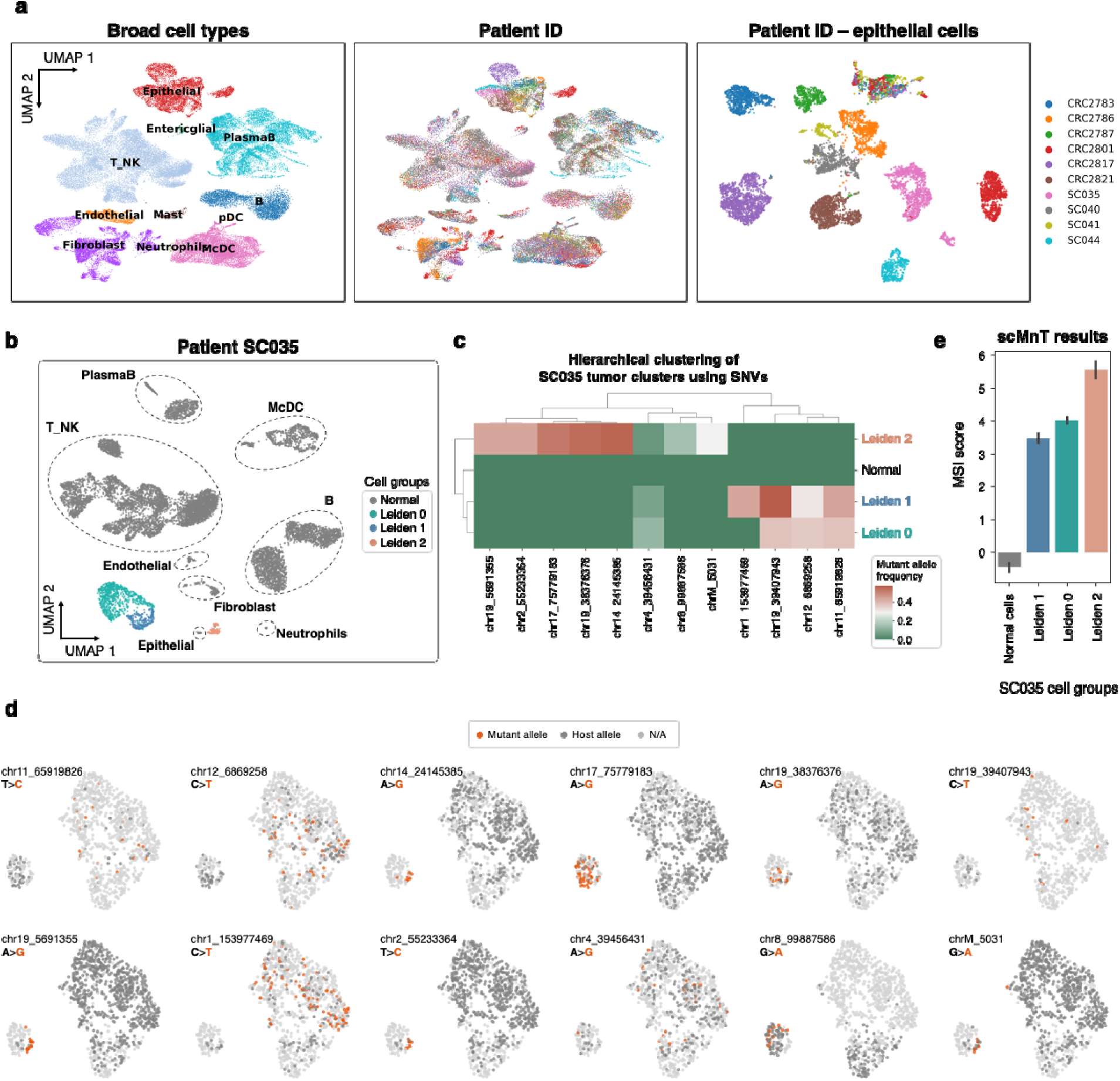
In-depth analysis of patient SC035 cancer cells reveals intra-tumoral heterogeneity in MSI intensity. **a**, UMAP visualization of all cells (left, middle) and epithelial cells (right) from the Joanito et al. (2022) dataset, colored by cell type (left) and patient ID (middle, right). **b**, UMAP visualization of cells from patient SC035 colored by their cell groups, defined as normal cells (gray) and cancer cells, with cancer cells further divided by their Leiden clusters. **c**, SNV-based hierarchical clustering of normal cells and the three cancer Leiden clusters from patient SC035. **d**, UMAP visualization of cancer cells from patient SC035 colored by their SNVs. **e**, Bar plot showing the MSI scores of each cell group from patient SC035.

**Extended Data Figure 3.**
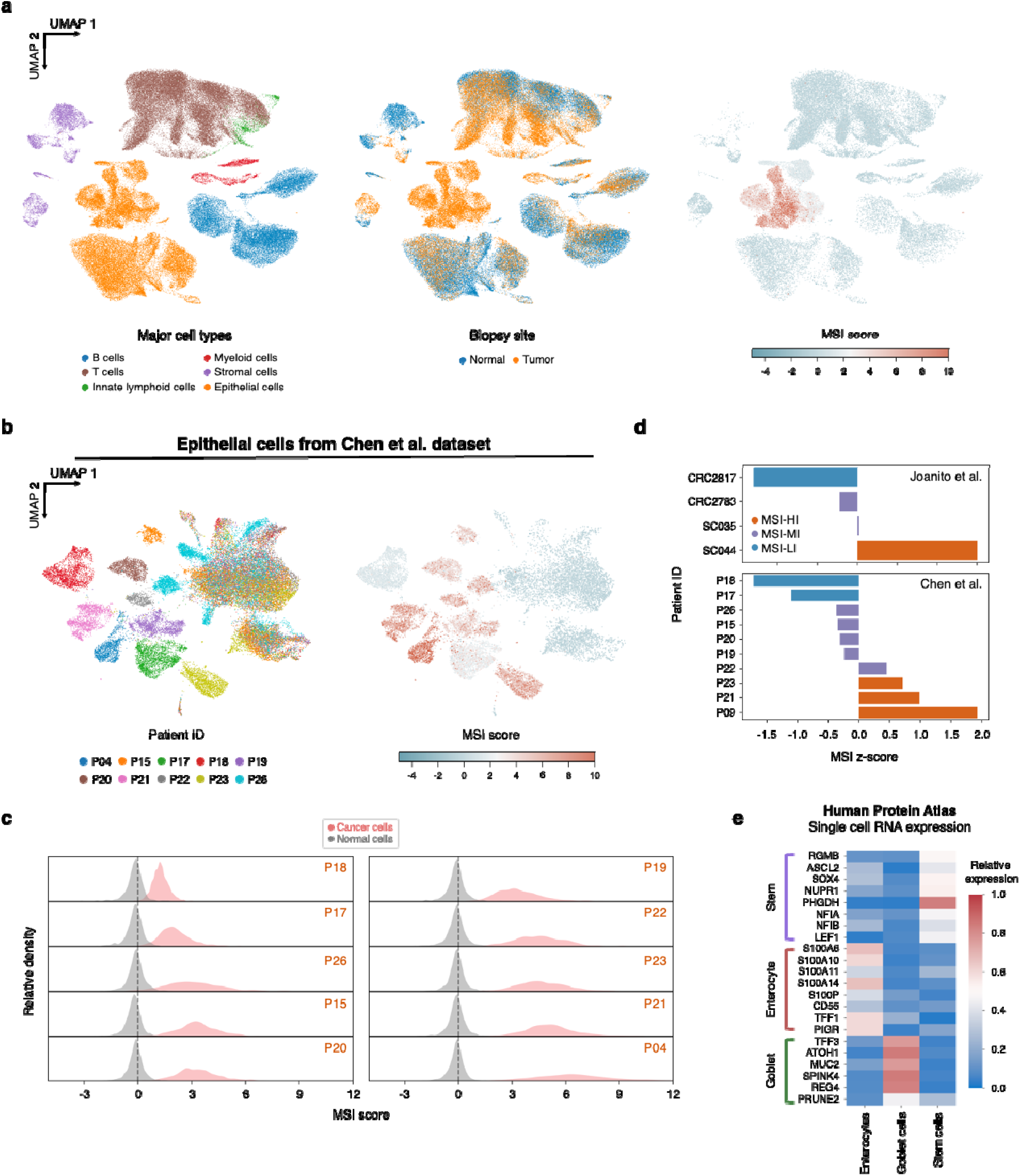
Application of scMnT in another colorectal cancer patient dataset from Chen et al. (2024). **a**, UMAP visualization of all cells from the Chen et al. (2024) dataset colored by major cell types (left), biopsy site (middle) and MSI score (right). **b**, UMAP visualization of cancer cells from the Chen et al. (2024) dataset colored by patient ID (left) and MSI score (right). **c**, KDE plot of MSI scores of normal cell (gray) and cancer cells (red) from the Chen et al. (2024) dataset. **d**, Standardized MSI scores of cancer cells within each dataset colored by the assigned MSI intensity group. **e**, Gene expression of selected MSI-HI and MSI-LI-associated DEGs in enterocytes, goblet cells, and stem cells from the Human Protein Atlas single cell type data.

